# Deviation from typical brain activity during naturalistic stimulation is related to personality traits

**DOI:** 10.1101/2024.04.23.586759

**Authors:** Lucia Jajcay, David Tomeček, Renáta Androvičová, Iveta Fajnerová, Filip Děchtěrenko, Jan Rydlo, Jaroslav Tintěra, Jiří Lukavský, Jiří Horáček, Jaroslav Hlinka

## Abstract

The relationship between personality and brain activity has been an increasingly popular topic of neuroscientific research. Naturalistic viewing has been shown to enhance individual differences and might, therefore, be particularly useful for exploring this relationship. Here, we thus examine neural signatures of personality using naturalistic fMRI of 82 healthy subjects. We implemented a simple dimensionality reduction method to characterize brain activity by its ‘typicality’, assessed a range of personality traits using widely-used personality inventories, and tested the relationship between the two. We found that there is, indeed, a relationship between personality and the typicality of brain activity, which appears to be most consistently manifested by lower typicality in subjects with higher Neuroticism/Harm Avoidance. Our results highlight the usefulness of naturalistic viewing data for exploring the relationship between individual differences in personality and brain activity.

## 1 INTRODUCTION

Personality refers to a set of cognitive, emotional, and behavioral patterns characteristic of an individual (American Psychological Association, 2018, April 19). These patterns are shaped by a combination of biological and environmental factors and tend to be relatively consistent and stable over time (Corr & Matthews, 2020). While there is no general agreement on a definition of personality, most definitions emphasize the role of motivation and interaction with the environment in shaping an individual’s unique behavioral and emotional responses to their surroundings, and revolve around personality traits.

Measuring personality is a complex endeavor that requires reliable and valid assessment tools. Personality questionnaires, or inventories, such as the NEO Five-Factor Inventory (NEO-FFI) (Costa Jr, 1992), which assesses the so-called Big Five personality traits (Digman, 1990), present one way of doing so (Corr & Matthews, 2020). They are standardized measures designed to assess the personality traits of an individual by asking them to rate themselves on a series of statements, and remain the dominant method of personality assessment (Corr & Matthews, 2020).

While traditional approaches to understanding personality have predominantly focused on behavioral and psychological perspectives, in recent years, there has been an increased interest in linking individual differences in personality to anatomical and functional brain markers.

Indeed, a growing amount of research has successfully detected relationships between personality measures and brain function. Due to its high spatial resolution, availability, and non-invasive nature, such ‘personality neuroscience’ studies often rely on functional magnetic resonance imaging (fMRI) (Dubois, Galdi, Han, Paul, & Adolphs, 2018; Tomeček et al., 2020).

Notably, several studies have used resting-state fMRI to relate brain functional connectivity (FC) to personality traits. FC captures statistical dependencies of the activity of spatially separated brain regions (Friston, 1994). It has been suggested that the FC matrix may be a useful representation of brain activity dynamics and provide behaviorally or clinically relevant markers (Biswal et al., 2010; van Dijk et al., 2010). FC has also been shown to act as a unique and reliable ‘fingerprint’ capable of distinguishing an individual by capturing their intrinsic, characteristic connectivity profile (Finn et al., 2015). Aghajani et al. (2014) found Neuroticism and Extraversion to be associated with different amygdala resting-state functional connectivity patterns. A study of the ‘neurotic brain’ by Servaas et al. (2015) concluded that the brains of highly neurotic individuals have less than optimal functional network organization and show signs of functional disconnectivity. Sampaio et al. (2014) revealed correlations between each of the Big Five personality dimensions and the default mode network (DMN). Similarly, Simon et al. (2020) found Openness to be positively associated with connectivity in the DMN. In addition, they found negative associations between Neuroticism and both the ventral and dorsal attention networks and Agreeableness and the dorsal attention network. They also showed that the association between Conscientiousness and the frontoparietal network becomes stronger with age. Moreover, the relationships between personality traits and resting-state functional connectivity also appear to be gender-specific, as demonstrated by Nostro et al. (2018). Wang et al. (2018) used an exploratory approach based on independent component analysis to identify a parietal network and found the inter-subject similarity of this network to be associated with Openness. Using connectome-based predictive modeling, Hsu et al. (2018) identified networks consisting of functional connections correlated with Neuroticism and Extraversion scores, then successfully predicted these scores for a different sample of individuals.

A more recent study by Cai et al. (2020) applied a similar approach to a large Human Connectome Project (HCP) dataset and could reliably predict four of the Big Five personality traits (all but Extraversion). Also capitalizing on the large HCP database, Liu et al. (2019) demonstrated that subjects with similar personality profiles have similar whole-brain connectivity patterns. Lastly, using individual FC matrices of 884 young, healthy adults, Dubois et al. (2018) reliably predicted Openness to experience. In addition, they also performed principal component analysis (PCA) on the personality scores to derive two orthogonal higher-order personality dimensions. They could reliably predict 5% of the variance in the score of the second component (β), which loaded mostly on Openness to experience. It, therefore, appears that personality can indeed be successfully investigated using fMRI. However, as argued by Tomeček et al. (2020), due to methodological challenges – arising from multiple testing and variability in analysis pipelines, among other things – the reflection of personality in the brain’s intrinsic functional architecture remains elusive, and findings of such studies should be interpreted with great caution.

Note that all of the above-mentioned studies relied on resting-state functional connectivity in conjunction with the five-factor model of personality (Digman, 1990). The naturalistic viewing (movie-watching) condition presents an increasingly popular alternative. Since it provides rich stimulation and, simultaneously, sufficient space for spontaneous responses, it might elicit highly individual responses and inter-regional interactions. Indeed, while naturalistic stimuli make individual activation profiles more similar to one another, observable individual differences have been shown to persist and even become easier to identify (Geerligs, Rubinov, Cam-CAN, & Henson, 2015; Vanderwal et al. 2017; Wang et al., 2017; Finn et al., 2017; Feilong, Nastase, Guntupalli, & Haxby, 2018; Finn & Bandettini, 2021).

Here, we therefore analyzed fMRI data obtained while subjects were watching movies. The more controlled nature of the naturalistic viewing condition allowed us to go beyond the FC approach commonly applied to resting-state data and focus on comparing the time courses of brain activations across subjects. Since the naturalistic viewing condition does not involve a structured task, standard general linear models are not directly applicable for modeling responses. To address this, we characterized brain activity during the condition by its ‘typicality’ – a straightforward, one-dimensional measure that quantifies how closely individual brain activity aligns with the group-average response to the stimulus, similarly to how beta-weights in standard task analyses capture alignment with the HRF-convolved stimulation paradigm. We then tested its relationship to personality traits, assessed using two well-established personality inventories. In addition to the widely-used NEO-FFI, we also used a second inventory based on a psychobiological model.

We expect that the naturalistic viewing condition combined with the comparably large dataset and comprehensive assessment of personality traits will allow us to examine the elusive relationship between personality and brain activity.

## 2 MATERIALS AND METHODS

### 2.1 Participants

Eighty-two healthy volunteers participated in the study (77 right-handed, 48 male, mean age ± SD: 30.96 ± 8.54 years). Participants were informed about the experimental procedures and provided written informed consent. The study design was approved by the local Ethics Committee of the Institute for Clinical and Experimental Medicine. Each participant underwent MRI scanning that included functional magnetic resonance imaging acquisitions during an audiovisual stimulation by segments of naturalistic stimuli (3 video segments, total length 29’45”) and 10 min in the resting-state condition with eyes closed, as well as the acquisition of a T1-weighted and T2-weighted anatomical scan, and a diffusion-weighted imaging scan.

### 2.2 Personality inventories

The participants filled out Czech versions of two widely-used personality inventories – the NEO-FFI (Costa Jr, 1992) and the Temperament and Character Inventory (TCI) (Cloninger, Przybeck, Svrakic, & Wetzel, 1994) – providing a total of 12 scores representing different personality traits.

The NEO-FFI consists of 60 items and measures five personality traits: Neuroticism, Extraversion, Openness to experience, Agreeableness, and Conscientiousness – the so-called Big Five. The five-factor model of personality is based on the lexical hypothesis, which assumes that fundamental individual differences are represented in the natural language by trait descriptive adjectives and was proposed as a universal and comprehensive framework for describing personality (Digman, 1990).

The TCI (240 items) is a seven-factor model of personality based on a psychobiological theory. It assesses four temperament dimensions assumed to be related to biology, heritable, and stable through the life-span: Novelty Seeking, Harm Avoidance, Reward Dependence, and Persistence; and three character dimensions affected by learning and culture, which are less heritable and mature with age: Self-Directedness, Cooperativeness, and Self-Transcendence (Aluja & Blanch, 2011; De Fruyt, Van de Wiele, & Van Heeringen, 2000).

Upon visual inspection, the distributions of most of the personality scores in our subject sample appear to be close to normal (see Fig. 1), yet a formal statistical test rejected normality for two of them (see Analysis).

**FIGURE 1.**
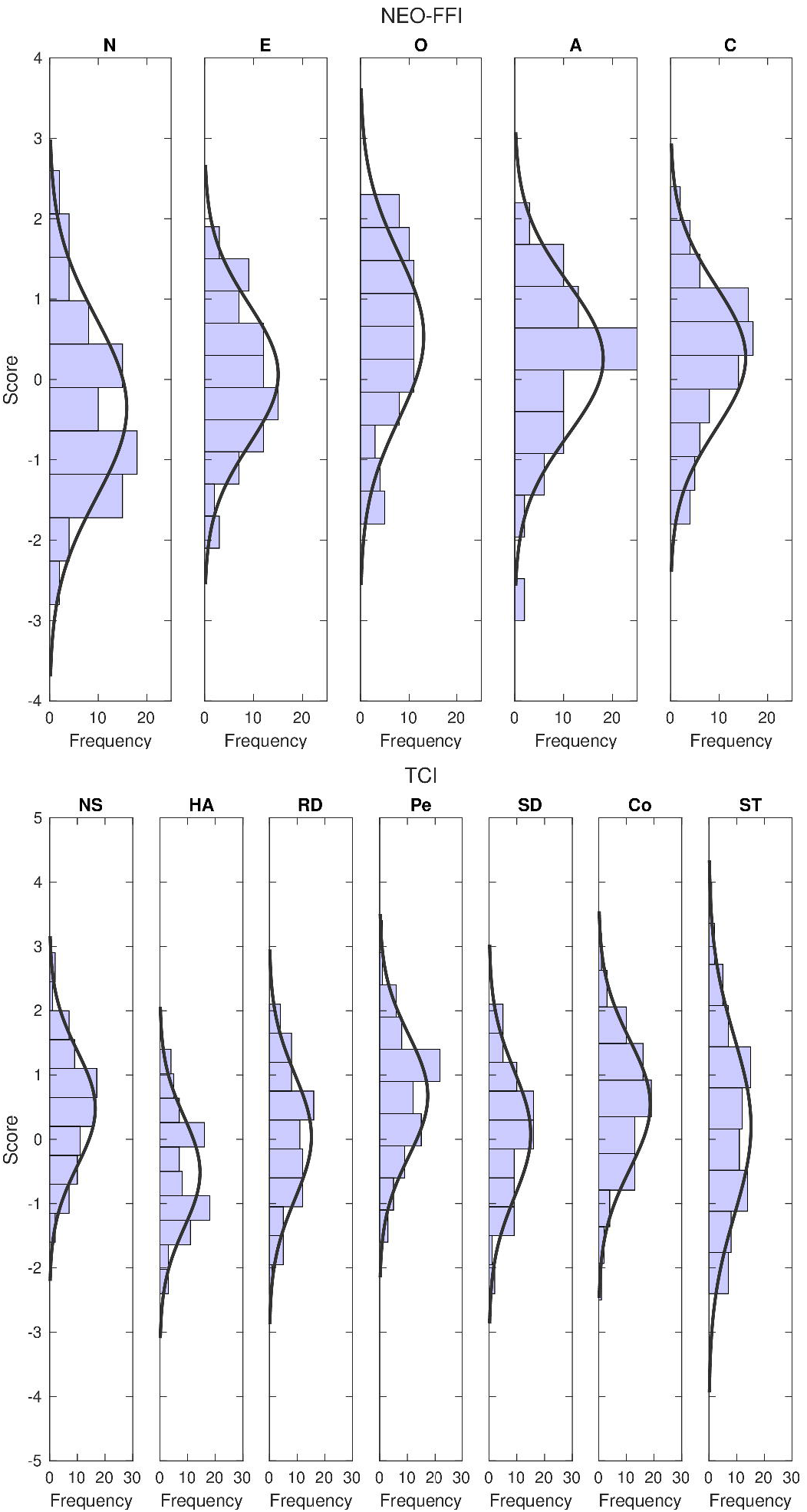
Distributions of the 12 personality traits in our subject sample (*N* = 82), with fitting normal distributions. *Top*: NEO-FFI scores; *bottom*: TCI scores.

Note that the Big Five are not fully orthogonal (Saucier, 2002). Similarly, certain TCI dimensions tend to correlate (Aluja & Blanch, 2011). Moreover, since the two inventories are measures of two competing theories of personality and might be measuring similar personality constructs with different scales, similarities and correlations naturally exist across the two. Most notably, Harm Avoidance (defined by Cloninger (1987, p. 575) as ‘a heritable tendency to respond intensely to signals of aversive stimuli, thereby learning to inhibit behavior to avoid punishment, novelty, and frustrative nonreward’) appears to be highly positively correlated with Neuroticism (the tendency to experience negative emotions) and negatively correlated with Extraversion (a measure of how sociable, outgoing and energetic a person is) (Costa & McCrae, 1985; De Fruyt et al., 2000).

As shown in Fig. 2, significant correlations across the 12 personality traits also exist in our sample. In particular, we found strong positive Pearson correlations (after Benjamini-Hochberg FDR correction) between Neuroticism and Harm Avoidance (*r* = 0.751, *q* < 0.001) and Agreeableness and Cooperativeness (*r* = 0.721, *q* < 0.001) and strong negative correlations between Extraversion and Harm Avoidance (*r* = – 0.590, *q* < 0.001) and Neuroticism and Self-Directedness (*r* = –0.585, *q* < 0.001). The relationships between the traits in our final sample are, therefore, in line with the findings of an analogous study mentioned above (De Fruyt et al., 2000).

**FIGURE 2.**
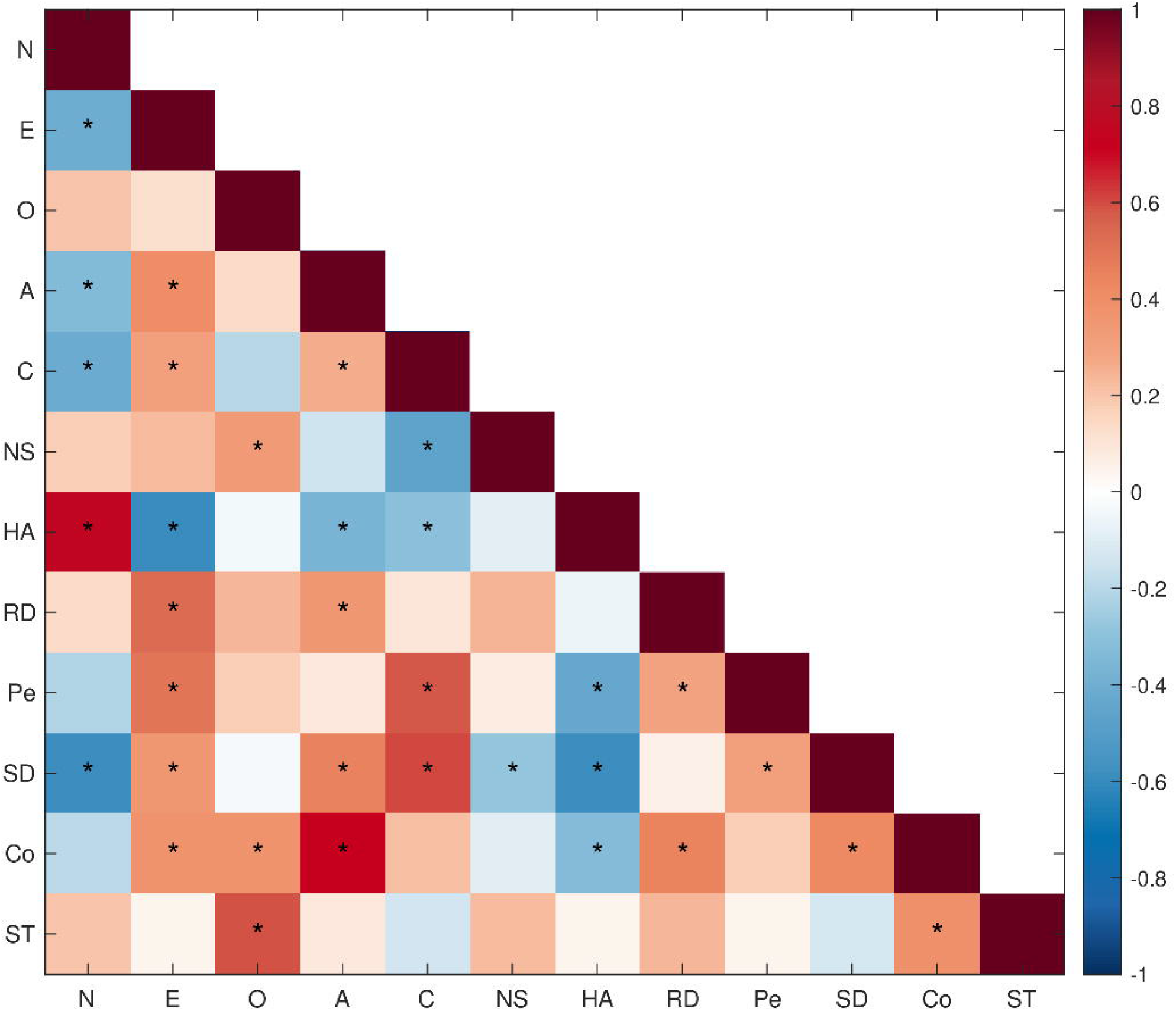
Relationships between the 12 personality traits in our subject sample (N = 82). Pearson correlations between traits assessed by NEO-FFI (N, E, O, A, C) and TCI (NS, HA, RD, Pe, SD, Co, ST). Asterisks indicate statistical significance after Benjamini-Hochberg FDR correction (q < 0.05).

### 2.3 Stimuli

We used 3 video segments, in the following order:

1. an uninterrupted excerpt from ‘The Good, the Bad and the Ugly’ (16:48-26:48), a feature film by Sergio Leone (1966) which has been used in previous studies of the naturalistic viewing condition (Hasson, Nir, Levy, Fuhrmann, & Malach, 2004; Mantini et al., 2012) (further denoted as ‘western’, duration 10 min);
2. an excerpt from a local TV reality show ‘Hledá se táta a máma’ (season 3, episode 2), which showed a strong activation profile in our previous, unpublished study (‘reality show’, 8 min and 5 s);
3. a collection of six short clips intended to elicit a range of emotions – excerpts from mostly Hollywood movies, from the FilmStim database (Schaefer, Nils, Sanchez, & Philippot, 2010) (‘emotional clips’, 11 min and 40 s).

Our selection of stimuli thus consists of materials high in social content and likely to elicit strong emotions and, similarly to other naturalistic viewing studies, includes well-known Hollywood movies (Finn & Bandettini, 2021; Vanderwal et al., 2017).

### 2.4 MRI data acquisition

Scanning was performed using a Siemens TrioTim 3T MRI machine located at the Institute for Clinical and Experimental Medicine in Prague, Czech Republic. A high-resolution T1-weighted image (MP-RAGE 3D, TR = 2300 ms, TE = 4.63 ms, TI = 900 ms, flip angle = 10°, voxel size = 1*×*1*×*1 mm, FOV 256*×*256, PAT-factor = 2, BW = 130 Hz/pix, 224 sagittal slices) covering the entire brain was acquired for anatomical reference. Functional images were then obtained using T2*-weighted echo-planar imaging (EPI) with blood oxygenation level-dependent (BOLD) contrast. GE-EPIs (TR = 2500 ms, TE = 30 ms, flip angle = 90°, voxel size = 3*×*3*×*3 mm, FOV 192*×*192, BW = 1594 Hz/pix, 44 axial slices) covering the entire cerebrum were acquired continuously in ascending order. T2-weighted and diffusion-weighted images were also acquired but not used in the current study.

The video segments were concatenated into a single MP4 file without breaks between the segments. They were back-projected onto a screen using an Epson EB-G5100 projector with an ELPLM04 (Middle Throw 1) lens. The subjects viewed the 47*×*35 cm large screen via a mirror mounted on the head coil at a 22° visual angle from a distance of 1210 mm. The audio was delivered using MRI-compatible headphones (MR CONFON OPTIME 1), with its intensity amplified, allowing the subjects to hear it over the scanner noise. For further technical details, such as the resolution and frame rate of each video, see Hlinka et al. (2022).

### 2.5 Preprocessing

Preprocessing was performed using the SPM12 software package (Wellcome Centre for Human Neuroimaging, UCL) implemented in MATLAB (The Mathworks, Inc.). The preprocessing pipeline consisted of motion correction, slice timing correction, spatial normalization into standard stereotaxic space (Montreal Neurological Institute, MNI), and spatial smoothing with an 8 mm FWHM kernel.

Subsequently, two types of time series to be used in further analyses were extracted from the data. Firstly, using the GIFT toolbox, Infomax-based independent component analysis (ICA) was performed, decomposing the data into 39 independent components. The number of components was estimated using the minimum description length (MDL) criterion with default settings. While removing artifactual components is, in many contexts, common practice, ICs are often a mixture of meaningful and noisy signal; manual labeling of artifacts is inherently subjective; and discarding components – regardless of the labeling method – can introduce bias (Griffanti et al., 2017; Salimi-Khorshidi et al., 2014). Here, we therefore retained all components, adopting a fully data-driven approach that preserves as much neural signal as possible, avoids labeling-related bias, and facilitates a reproducible analysis. The signal was band-pass filtered using the Butterworth filter (0.009–0.1 Hz).

Secondly – for an alternative, region-based processing pipeline – mean BOLD time series from the 90 regions of interest (ROIs) of the Automated Anatomical Labeling (AAL) atlas were extracted. The anatomical positions of the ROIs are described in Tzourio-Mazoyer et al. (2002). The signal was band-pass filtered using the same Butterworth filter (0.009–0.1 Hz).

### 2.6 Analysis

#### 2.6.1 ICA analyses

We first calculated the ‘typical time series’ (average across all subjects) for each of the 39 independent components. For each subject, we then correlated each of their time series with the respective typical time series. The mean of these Pearson’s correlation coefficients (39 ‘similarity indices’) presents the subject’s individual ‘typicality index’ – a single number characterizing the typicality of their brain activity.

Using the interquartile range method, selected for its robustness, we identified three subjects with low typicality indices as outliers.

Subjects were considered outliers if their typicality indices were more than 1.5 interquartile ranges above the upper quartile (75th percentile) or below the lower quartile (25th percentile). To prevent extreme values from skewing the results, we removed these subjects and then repeated the analysis to obtain the final typicality indices. We thus ultimately evaluated typicality on a slightly reduced but robust dataset.

Since demographic factors such as age and sex have been reported to be related to the relationship between personality traits and brain activity, we examined the relationship between the typicality indices and these two potential confounds before proceeding with the analyses (Nostro et al., 2018; Simon et al., 2020). While the typicality indices are normally distributed, the null hypothesis of a normal distribution was rejected for subject age and sex (using the Anderson-Darling test implemented in MATLAB; *p* < 0.05). We thus assessed the relationship between the typicality indices and these two variables by computing Spearman’s rank correlations.

As discussed above, similarities and overlaps across the traits assessed by the two personality inventories have been shown to exist in the general population as well as in our sample. To examine the relationship between the typicality indices and personality traits, we thus reduced the dimensionality of the personality data by performing PCA. In our primary analysis, we calculated Spearman’s rank correlation between the typicality indices and a subset of the principal components. The decision to use *Spearman’s rank* correlation was made because, using the Anderson-Darling test implemented in MATLAB, the null hypothesis of a normal distribution was rejected for Agreeableness (NEO-FFI) and Harm Avoidance (TCI). We performed the analysis for all stimuli combined (using the entire time series corresponding to all three videos) as well as for individual stimuli (using the corresponding segment of the entire time series) to assess robustness with respect to stimulus selection. All *p*-values were corrected for multiple comparisons using the Benjamini-Hochberg FDR procedure; the resulting *q*-values are reported. We then performed a number of exploratory follow-up analyses. The same FDR correction was applied to these to ensure consistent control of false positives.

First, after observing a significant relationship between the high-level (PCA-based) personality traits and the typicality of brain activity, we explored the result in further detail by calculating Spearman’s rank correlation between subjects’ typicality indices and their scores on each of the twelve, albeit overlapping, personality traits. Again, we performed this analysis for all stimuli combined as well as for individual stimuli.

Next, we delved into even more detail and examined the relationship between each ICA component and the personality traits – meaning we computed Spearman’s rank correlation between the latter and the 39 correlation coefficients of each subject’s time series with the respective typical time series (their ‘similarity indices’ rather than their summarized ‘typicality index’).

The doubly multivariate nature of the data (with multiplicity in terms of both personality traits and ICA components) invites the application of a more advanced technique. In addition to simply computing pairwise correlations, we thus also performed a canonical correlation analysis (CCA; Krzanowski & Krzanowski, 1988; Seber, 1984). The latter is an alternative method of measuring the association among the two sets of variables – the 12 personality scores on the one hand and the 39 similarity indices on the other. It finds linear combinations of variables from the two sets (canonical variables) with maximum correlations. Note that this analysis was not corrected for multiple comparisons, as it represents a single multivariate test rather than multiple pairwise tests. It is included for the sake of completeness and, while advanced and appropriate, the large number of variables and the relatively small sample size mean that even CCA is unlikely to yield statistically significant results.

#### 2.6.2 AAL analyses

We repeated the analyses using the mean BOLD time series from the 90 ROIs derived from the AAL atlas rather than the ICA components. The canonical correlation analysis could not be performed on the AAL data due to rank deficiency, resulting from having more features (90 time series) than observations (81 subjects). Note that only one of the three outliers removed in the ICA analysis was removed in the AAL analysis. The same IQR criterion was used in both cases.

## 3 RESULTS

### 3.1 ICA analyses

Since neither of the correlations between subject typicality indices and subject age or sex is statistically significant (age: *ρ* = –0.139, *p* = 0.220; sex: *ρ* = –0.120, *p* = 0.293), the following results do not take age or sex into account.

The principal component analysis performed on the personality data produced 12 principal components. Based on Fig. 3, the first two principal components capture most of the relevant between-subject variability. We thus computed Spearman’s rank correlation between the typicality indices and the *first two* principal components. Note that Fig. 4 shows that the first principal component is influenced negatively by Neuroticism and Harm Avoidance and positively primarily by Cooperativeness and Agreeableness – the two pairs of personality traits shown above to be the most strongly correlated among the 12 traits.

**FIGURE 3.**
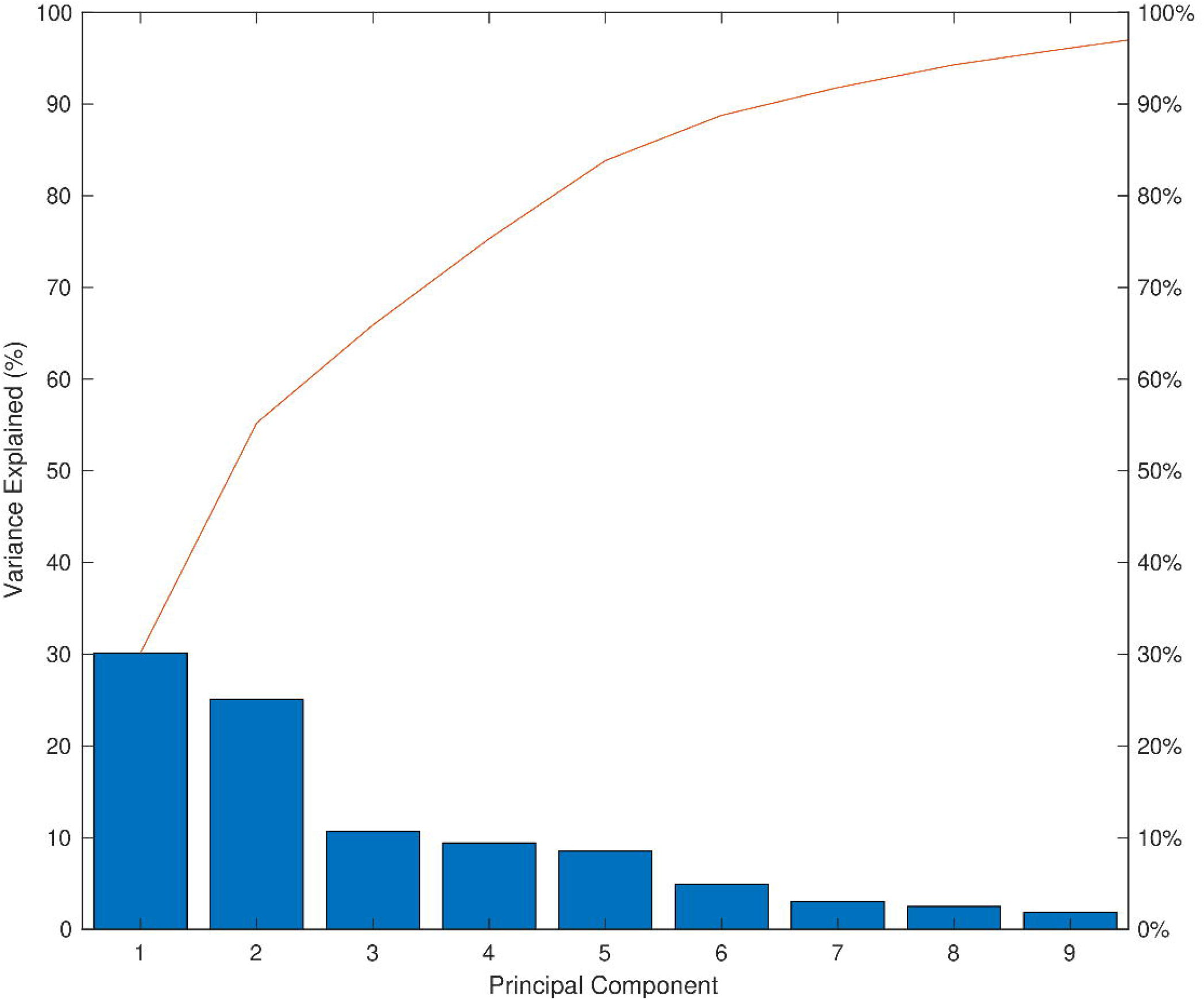
Percentage of total variance explained by the principal components (ICA analysis). The bars show the percentage of variance explained by each component. The line plot shows the cumulative explained variance percentages. The plot only shows the first nine components that together explain 95% of the total variance. A bend in the line occurs after the first two components, which together explain more than half (55%) of the total variance. Only the first two components were therefore used in the subsequent analysis.

**FIGURE 4.**
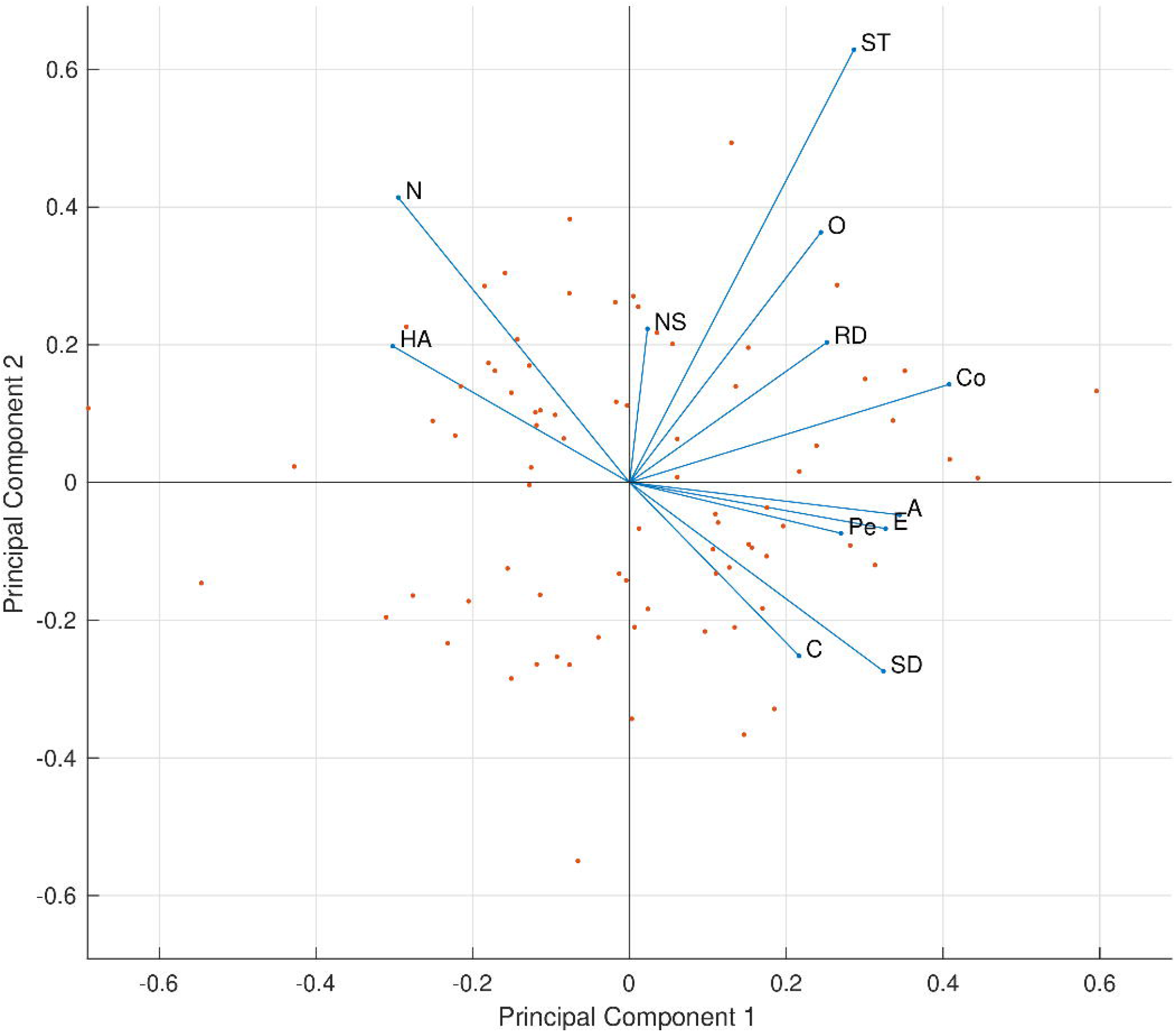
PCA biplot showing the influence of each personality trait on the principal components (ICA analysis). The 12 variables (personality traits) are depicted as vectors with their direction and length indicating how each trait contributes to the first two principal components (depicted on the x-axis and the y-axis, respectively) – principal component *coefficients* or loadings. In addition, the 79 observations (participants) are represented as points with their coordinates indicating each participant’s *scores* for the two principal components. The points are scaled with respect to the maximum score value and maximum coefficient length.

We found the typicality of brain activity during the experimental condition to be positively correlated with the first principal component, which represents the maximum variance direction of the data along the personality axis (*ρ* = 0.324, *q* = 0.007). The negative correlation with the second principal component is not statistically significant (*ρ* = – 0.171, *q* = 0.131). Similarly, when considering individual video segments, only the correlation with the first principal component is statistically significant, for the reality show (1st PC: western: *ρ* = 0.230, *q* = 0.084; reality show: *ρ* = 0.363, *q* = 0.007; emotional clips: *ρ* = 0.254, *q* = 0.072. 2nd PC: western: *ρ* = –0.154, *q* = 0.195; reality show: *ρ* = –0.147, *q* = 0.195; emotional clips: *ρ* = –0.158, *q* = 0.195). For comparison and full transparency, the Appendix also includes the results of this analysis without removing outliers (Table A1) and with age and sex included as covariates (Table A3).

Upon closer inspection of individual personality traits, we found that the typicality of brain activity during the experimental condition is significantly negatively correlated with Neuroticism (*ρ* = –0.290, *q* = 0.048) and Harm Avoidance (*ρ* = –0.307, *q* = 0.048) and positively correlated with Cooperativeness (*ρ* = 0.281, *q* = 0.048). These relationships are depicted in Fig. 5. No significant relationship with the other personality traits was observed. Looking at the individual video segments, the brain activity corresponding to the reality show shows a significant positive correlation with Extraversion (*ρ* = 0.360, *q* = 0.040). The remaining results are descriptively similar to the pattern observed across all stimuli, albeit not statistically significant. For detailed results, see Table 1.

**TABLE 1.** Relationship between each personality trait and the typicality of brain activity for individual video segments (ICA analysis). Spearman’s rho (*q*-value) for each video is shown. Results across all videos (‘all’) are also included for comparison. Statistically significant (*q* < 0.05, FDR-corrected) results are highlighted.

|  |  | all | western | reality show | emotional clips |
| --- | --- | --- | --- | --- | --- |
| NEO-FFI | Neuroticism | **-0.290 (0.048)** | -0.243 (0.156) | -0.290 (0.087) | -0.201 (0.205) |
|  | Extraversion | 0.265 (0.055) | 0.233 (0.156) | **0.360 (0.040)** | 0.143 (0.367) |
|  | Openness to experience | 0.003 (0.980) | -0.091 (0.639) | 0.053 (0.769) | -0.033 (0.867) |
|  | Agreeableness | 0.169 (0.233) | 0.107 (0.570) | 0.201 (0.205) | 0.141 (0.367) |
|  | Conscientiousness | 0.149 (0.284) | 0.073 (0.709) | 0.181 (0.247) | 0.150 (0.360) |
| TCI | Novelty Seeking | 0.046 (0.822) | 0.062 (0.731) | 0.068 (0.709) | 0.001 (0.996) |
|  | Harm Avoidance | **-0.307 (0.048)** | -0.277 (0.097) | -0.304 (0.078) | -0.192 (0.215) |
|  | Reward Dependence | 0.022 (0.924) | 0.006 (0.996) | 0.077 (0.709) | 0.003 (0.996) |
|  | Persistence | 0.216 (0.113) | 0.149 (0.360) | 0.218 (0.193) | 0.213 (0.194) |
|  | Self-Directedness | 0.215 (0.113) | 0.100 (0.592) | 0.264 (0.112) | 0.169 (0.290) |
|  | Cooperativeness | **0.281 (0.048)** | 0.198 (0.205) | 0.315 (0.078) | 0.233 (0.156) |
|  | Self-Transcendence | -0.049 (0.822) | -0.044 (0.816) | -0.030 (0.867) | -0.071 (0.709) |

**FIGURE 5.**
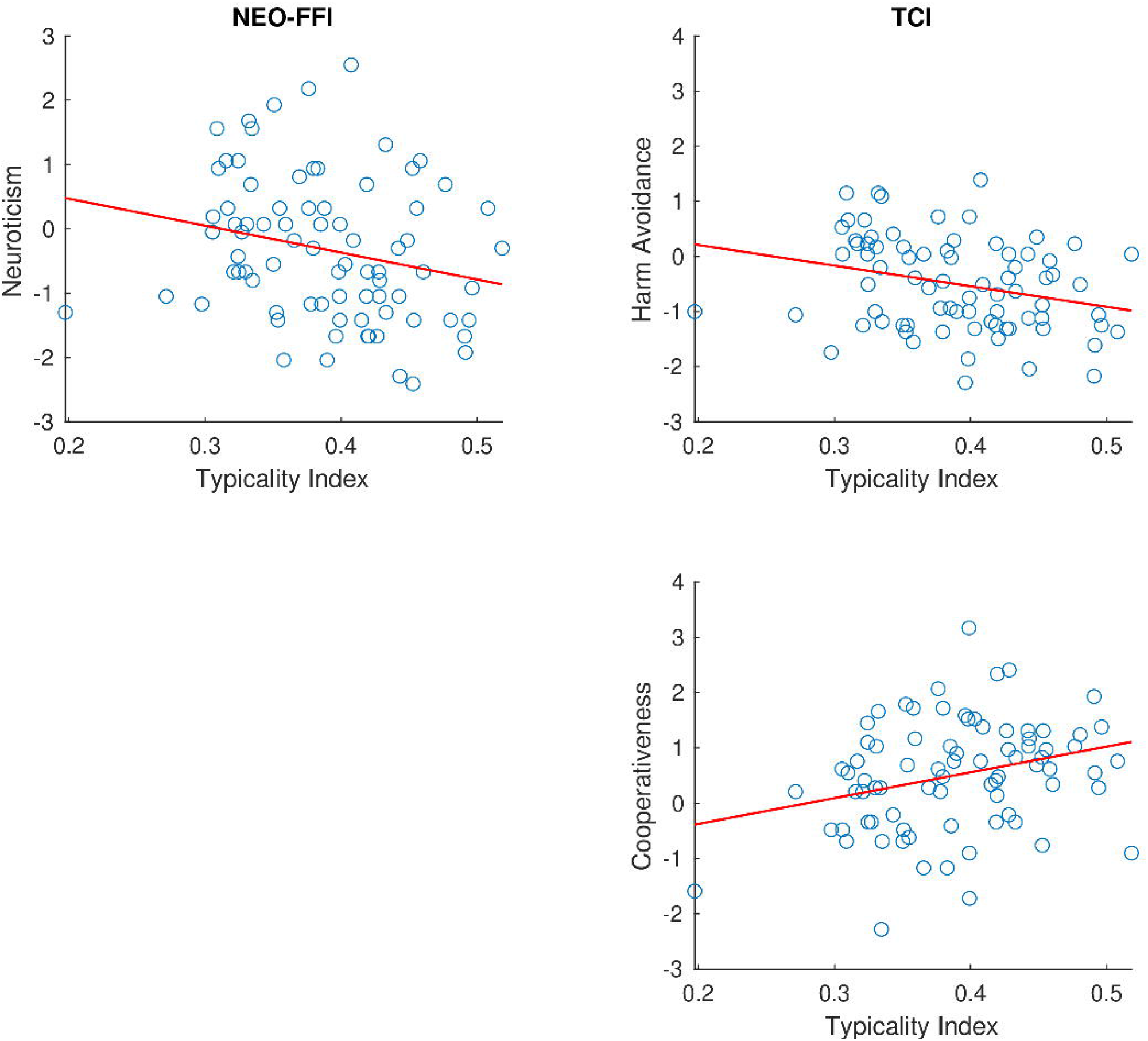
Relationship between the typicality of brain activity measured across the three video segments and selected personality traits (ICA analysis). The scatterplots show relationships between the typicality indices (horizontal axes) and the scores on certain personality dimensions of the two personality inventories (vertical axes) across subjects. Only personality traits for which the result is statistically significant (q < 0.05, FDR-corrected) are shown. Left (NEO-FFI): top: Neuroticism (ρ = –0.290, q = 0.048). Right (TCI): top: Harm Avoidance (ρ = –0.307, q = 0.048); bottom: Cooperativeness (ρ = 0.281, q = 0.048).

Figure 6 shows how each of the 39 ICA components relates to each of the 12 personality traits. Note that the pattern is primarily driven by the personality traits rather than by the similarity indices corresponding to the independent components of brain activity. While none of the correlations survived FDR correction, the general directions of the correlations observed with uncorrected *p*-values are consistent with the patterns reported for the summarized typicality index: all significant correlations with Neuroticism and Harm Avoidance are negative, while those with Extraversion and Cooperativeness are positive. The values are, naturally, affected by statistical power being weaker in the case of individual similarity indices relative to the summarized typicality index. The canonical correlation analysis resulted in pairs of canonical variables with correlations ranging from *r* = 0.428 to *r* = 0.911. Notably, none of the canonical correlations are statistically significant (strongest: *F*(468,364) = 1.110, *p* = 0.146; all others: *p* > 0.05). This lack of significance, despite relatively high observed *r*-values, is due to the unfavorable ratio of the number of variables to the number of samples (subjects) in the analysis.

**FIGURE 6.**
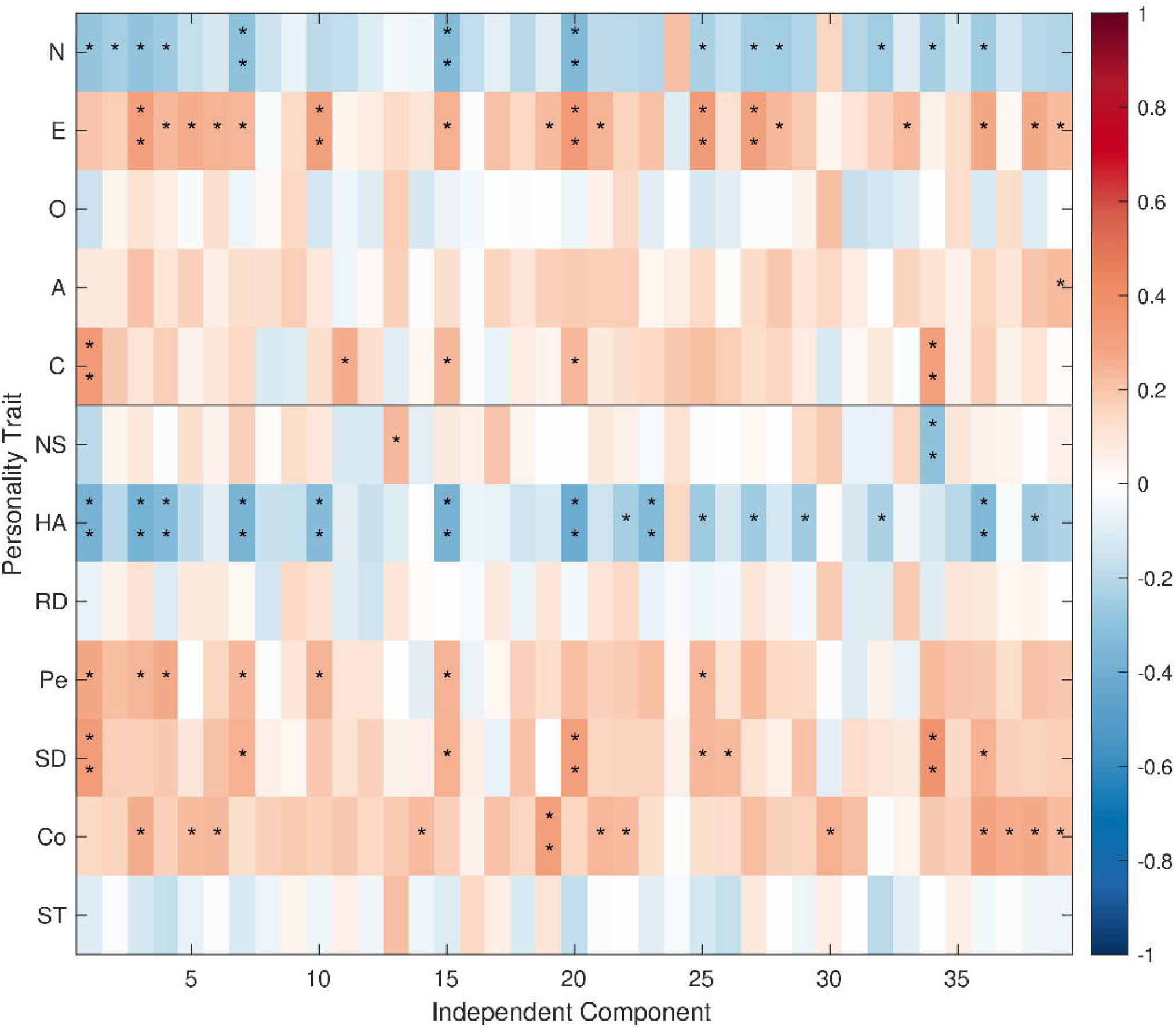
Spearman correlations between the 12 personality traits and the similarity indices corresponding to the 39 independent components (ICA analysis). Asterisks indicate uncorrected p-values (* p < 0.05, ** p < 0.01); no correlations survive FDR correction.

### 3.2 AAL analyses

Note that, due to the removal of different subsets of outliers, the final subject sample for the AAL analyses (*N* = 81) is slightly larger than that used in the ICA analyses (*N* = 79). Nevertheless, neither of the correlations between the typicality indices and subject age and sex is statistically significant (age: *ρ* = –0.181, *p* = 0.106; sex: *ρ* = –0.018, *p* = 0.871). The following analyses, thus, again, do not take these two variables into account.

The first two principal components, again, explain 55% of the total variance and were thus selected for the primary analysis. Here, the first principal component is influenced negatively by Neuroticism and Harm Avoidance and positively primarily by Cooperativeness and Self-Directedness, i.e., with this data, the latter surpasses Agreeableness.

Using the 90 regions of the AAL atlas instead of the 39 ICA components, the experimental condition is, again, positively correlated with the first principal component (*ρ* = 0.276, *q* = 0.026) and the negative correlation with the second principal component, again, is not statistically significant (*ρ* = –0.092, *q* = 0.413). Similarly, when considering individual video segments, only the correlation with the first principal component is statistically significant – and only for the reality show (1st PC: western: *ρ* = 0.193, *q* = 0.253; reality show: *ρ* = 0.297, *q* = 0.043; emotional clips: *ρ* = 0.154, *q* = 0.339. 2nd PC: western: *ρ* = – 0.069, *q* = 0.645; reality show: *ρ* = –0.046, *q* = 0.685; emotional clips: *ρ* = –0.081, *q* = 0.645). Again, the Appendixalsoincludestheresultsof this analysis without removing outliers (Table A2) and with age and sex included as covariates (Table A4).

The examination of individual personality traits showed no significant correlations. For details, see Table 2. While the results appear to be broadly consistent with the patterns observed in the ICA analysis (Table 1), given the lack of statistical significance, these apparent similarities should not be overinterpreted.

**TABLE 2.** Relationship between each personality trait and the typicality of brain activity for individual video segments (AAL analysis). Presentation as in Table 1.

|  |  | **all** | **western** | **reality show** | **emotional clips** |
| --- | --- | --- | --- | --- | --- |
| **NEO-FFI** | **Neuroticism** | -0.226 (0.158) | -0.195 (0.427) | -0.262 (0.130) | -0.086 (0.888) |
|  | **Extraversion** | 0.183 (0.246) | 0.141 (0.600) | 0.275 (0.116) | 0.012 (0.958) |
|  | **Openness to experience** | 0.045 (0.832) | -0.006 (0.958) | 0.116 (0.685) | -0.023 (0.950) |
|  | **Agreeableness** | 0.103 (0.620) | 0.065 (0.888) | 0.083 (0.888) | 0.081 (0.888) |
|  | **Conscientiousness** | 0.081 (0.630) | 0.022 (0.950) | 0.139 (0.600) | 0.054 (0.898) |
| **TCI** | **Novelty Seeking** | 0.086 (0.630) | 0.176 (0.427) | 0.068 (0.888) | -0.071 (0.888) |
|  | **Harm Avoidance** | -0.250 (0.150) | -0.288 (0.116) | -0.299 (0.116) | -0.006 (0.958) |
|  | **Reward Dependence** | 0.000 (0.997) | -0.056 (0.898) | 0.073 (0.888) | -0.009 (0.958) |
|  | **Persistence** | 0.132 (0.479) | 0.051 (0.898) | 0.184 (0.427) | 0.047 (0.898) |
|  | **Self-Directedness** | 0.249 (0.150) | 0.175 (0.427) | 0.278 (0.116) | 0.124 (0.657) |
|  | **Cooperativeness** | 0.216 (0.158) | 0.123 (0.657) | 0.179 (0.427) | 0.166 (0.457) |
|  | **Self-Transcendence** | -0.023 (0.913) | 0.040 (0.898) | -0.022 (0.950) | -0.040 (0.898) |

None of the correlations between the 90 ROIs and the 12 personality traits survived FDR correction. Figure 7 shows the general directions of the correlations, with significance indicated based on uncorrected *p-* values. At a glance, the patterns, again, appear to resemble those of the analogous ICA result. However, these are included purely for illustration and should be interpreted with caution.

**FIGURE 7.**
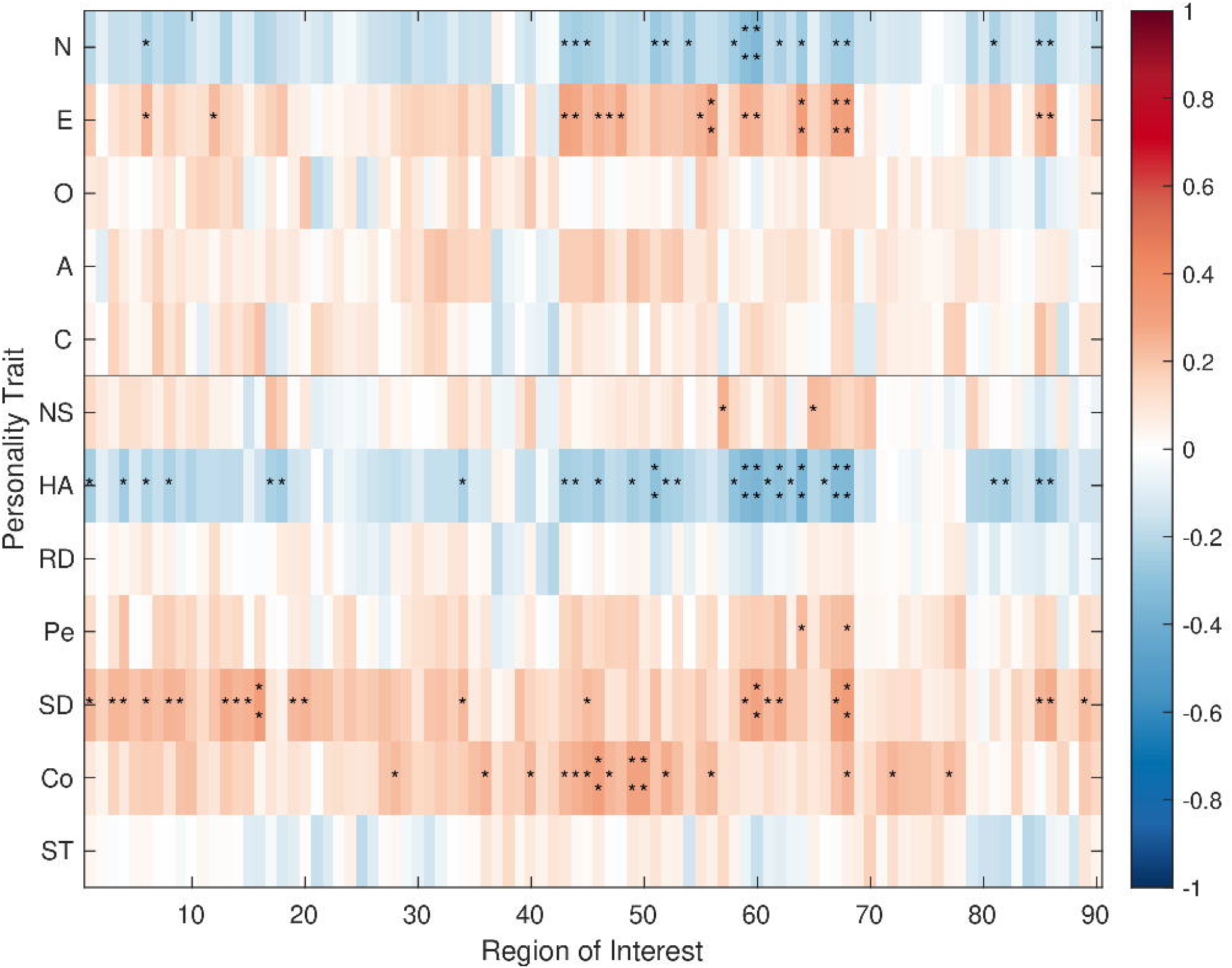
Spearman correlations between the 12 personality traits and the similarity indices corresponding to the 90 regions of interest (AAL analysis). Asterisks indicate uncorrected p-values (* p < 0.05, ** p < 0.01); no correlations survive FDR correction.

## 4 DISCUSSION

Using naturalistic fMRI thus helped us detect relationships between brain activity and personality. While the personality traits associated with the typicality of brain activity vary slightly across the analyses, the relationship between lower typicality and Neuroticism/Harm Avoidance – two highly correlated personality traits that likely measure similar aspects of personality – emerges across all of them, including i n the primary analysis where the first principal component loads mainly on these traits. Moreover, the ICA analyses suggest a positive relationship with Extraversion – perhaps unsurprisingly, given that Extraversion and Neuroticism are strongly negatively correlated.

The results of our primary analysis reveal a relationship between personality and the typicality of brain activity, detectable with ICA as well as AAL data. While this qualitative replication across the two methods does not constitute a completely independent validation of the results, as these are, in the end, just different but related summary statistics derived from the same brain activity, it provides some evidence of robustness with respect to methodological choices.

Comparisons across the video segments suggest that the relationship with personality traits slightly varies across naturalistic stimuli. However, we did not explicitly test for differences. Moreover, since the videos may differ in a range of parameters, we do not speculate on the generalizability of the observations to entire genres. We note, nevertheless, that the results of these comparisons are in line with previous reports of social clips predicting behavior better than other types of videos (Finn & Bandettini, 2021). Moreover, the advantage of the reality show could also be due to its comparably high split-half reliability (Hlinka et al., 2022).

We further narrowed the results down to individual personality traits and related those to individual independent components/ROIs. None of the relationships appear to be significant after FDR correction. Uncorrected results should be interpreted cautiously. We present them for context and refrain from drawing conclusions about the role of different ICs or ROIs in relation to individual personality traits.

The more stringent canonical correlation analysis did not detect a relationship between personality traits and similarities of brain activations to those of an average subject. Given the large number of degrees of freedom – stemming from the large number of independent variables on both sides (personality traits and typicality indices) and the relatively small sample size – the fact that canonical correlation analysis lacked the power to detect the relationship and failed to replicate the previous findings is in line with expectations, as noted in the Analysis section.

While the main findings remained largely similar when time series derived from regions of the AAL atlas were used instead of independent components, statistically significant correlations were generally fewer and weaker. Including all anatomic regions of the brain will inevitably introduce more noise than the output of independent component analysis. The weaker correlations are, therefore, unsurprising and highlight the appropriateness of ICA for our study.

As explained above, the typicality index of a subject was computed by correlating their time series to the typical time series. The fact that a typical time series was computed as the mean across all subjects, including the subject in question, could arguably have had an impact on the results. While it clearly biases typicality upwards, the case for it also systematically affecting the relationship between personality and typicality is rather unclear. Still, out of an abundance of caution, we also performed a version of the analyses where a subject was omitted from the calculation of the typical time series when their typicality index was being quantified. As expected, the impact of the inclusion or removal of the subject on the result was negligible.

Since the unconstrained resting-state condition does not allow for meaningful comparisons of time series across subjects, previous studies relating personality to brain activity all analyzed functional connectivity. For comparison, we also repeated our primary analysis using functional connectivity – rather than computing the typicality of the time series, here, we computed the typicality of the FC matrices derived from them (both ICA and AAL). Interestingly, using this approach, we found subject age to be (negatively) correlated with subject typicality indices. This finding is in line with the previously reported age-related differences discussed above. The association between personality and FC typicality was therefore measured using Spearman’s *partial* correlation, controlling for age. We did not detect any statistically significant results. This highlights the importance of temporal dynamics and, consequently, the advantage of the naturalistic viewing condition for studies of the relationship between personality and brain activity.

## 5 CONCLUSION

We have shown that a relationship between personality traits and the typicality of brain activity, indeed, exists, and can be detected using naturalistic stimuli. While a number of personality traits seem to be related to the typicality of brain activity, this relationship is most evident in subjects with higher Neuroticism and Harm Avoidance whose brain activity tended to show lower typicality.

Our findings highlight the importance of temporal dynamics for studies of the relationship between personality and brain activity. Consequently, the fact that the naturalistic viewing condition allows for taking temporal dynamics into account presents an important advantage over rest.

The use of the newly defined single overall typicality index, as well as only a couple of main personality features for the primary analysis, provided a highly powerful statistical tool for evaluating the relationship between personality and brain activity in the naturalistic viewing condition. However, further research is necessary to enhance our understanding of the nature of this relationship. While our dataset is, indeed, relatively large and comprehensive, even larger datasets with a greater range of personality scores might further increase signal variance, thus increasing the likelihood of detecting associations. Validating the results using different, preferably larger, datasets could also allow for a more comprehensive examination of the relationship between personality traits and brain activation patterns, and more confidence in drawing conclusions regarding specific personality traits, as well as an improved understanding of the variability across different audiovisual stimuli. Deliberately choosing stimuli that differ with respect to a certain parameter, while being similar in other aspects, might provide more insights into the role of various parameters of the stimuli. This would, in turn, allow for better informed selection of suitable stimuli in future experiments.

## AUTHOR CONTRIBUTIONS

**Lucia Jajcay:** Methodology, Software, Validation, Formal analysis, Investigation, Writing - original draft, Writing - review & editing, Visualization. **David Tomeček:** Software, Formal analysis, Investigation, Writing - review & editing. **Renáta Androvičová:** Investigation, Data curation, Writing - review & editing, Project administration. **Iveta Fajnerová:** Investigation, Data curation, Writing - review & editing, Project administration. **Filip Dechtěrenko:** Investigation, Data curation, Writing - review & editing, Project administration. **Jan Rydlo:** Investigation, Writing - review & editing. **Jaroslav Tintěra:** Investigation, Resources, Writing - review & editing, Project administration, Funding acquisition. **Jiří Lukavský:** Investigation, Resources, Writing - review & editing, Supervision, Project administration, Funding acquisition. **Jiří Horáček:** Investigation, Resources, Writing - review & editing, Supervision, Project administration, Funding acquisition. **Jaroslav Hlinka:** Conceptualization, Methodology, Validation, Formal analysis, Investigation, Resources, Data curation, Writing - review & editing, Supervision, Project administration, Funding acquisition.

## FUNDING INFORMATION

This work was supported by the European Regional Development Fund (ERDF) – project Brain dynamics (project No. CZ.02.01.01/00/22_008/0004643); the Czech Science Foundation (project No. 21-32608S); the Johannes Amos Comenius Programme (P JAC) provided by MSMT (project No. CZ.02.01.01/00/23_025/0008715); and by the Grant Agency of the Czech Technical University in Prague (grant No. SGS23/119/OHK3/2T/13).

## CONFLICT OF INTEREST STATEMENT

The authors declare no conflict of interest.

## DATA AVAILABILITY STATEMENT

The fMRI and personality data are stored in a public OSF repository at https://osf.io/795hb/.

## SUPPORTING INFORMATION

Additional supporting information can be found in the online version of the article at the publisher’s website.

## Notes

Funding Information This research was supported by the European Regional Development Fund (ERDF) – project Brain dynamics (project No. CZ.02.01.01/00/22_008/0004643); the Czech Science Foundation (project No. 21-32608S); the Johannes Amos Comenius Programme (P JAC) provided by MSMT (projectNo. CZ.02.01.01/00/23_025/0008715); andbythe Grant Agency of the Czech Technical University in Prague (grant No. SGS23/119/OHK3/2T/13).

https://osf.io/795hb/

### Competing Interest Statement

The authors have declared no competing interest.

### Summary of Updates

Minor clarifications made in response to reviewer feedback.

