## Appendix for "Deviation from typical brain activity during naturalistic stimulation is related to personality traits"

TABLE A1

Main results with outliers removed (as reported in the manuscript) and not removed (for comparison). ICA analysis. Statistically significant results ($q < 0.05$, FDR-corrected) are highlighted. (See Note on component orientation at the end of the Appendix).

| outliers | PC | all | western | reality show | emotional clips |
|---|---|---|---|---|---|
| removed | 1st | **0.324 (0.007)** | 0.230 (0.084) | **0.363 (0.007)** | 0.254 (0.072) |
| | 2nd | −0.171 (0.131) | −0.154 (0.195) | −0.147 (0.195) | −0.158 (0.195) |
| not removed | 1st | **0.251 (0.047)** | 0.161 (0.295) | **0.302 (0.036)** | 0.173 (0.295) |
| | 2nd | −0.010 (0.926) | 0.010 (0.928) | −0.049 (0.928) | 0.014 (0.928) |

TABLE A2

Main results with outliers removed (as reported in the manuscript) and not removed (for comparison). AAL analysis. Statistically significant results ($q < 0.05$, FDR-corrected) are highlighted. (See Note on component orientation at the end of the Appendix).

| outliers | PC | all | western | reality show | emotional clips |
|---|---|---|---|---|---|
| removed | 1st | **0.276 (0.026)** | 0.193 (0.253) | **0.297 (0.043)** | 0.154 (0.339) |
| | 2nd | −0.092 (0.413) | −0.069 (0.645) | −0.046 (0.685) | −0.081 (0.645) |
| not removed | 1st | 0.231 (0.074) | 0.151 (0.527) | 0.270 (0.085) | 0.106 (0.685) |
| | 2nd | 0 (0.997) | −0.007 (0.953) | −0.037 (0.944) | 0.030 (0.944) |

TABLE A3

Main results with age and sex not included as covariates (as reported in the manuscript) and with age and sex included as covariates (for comparison). ICA analysis. Statistically significant results ($q < 0.05$, FDR-corrected) are highlighted.

| covariates | PC | all | western | reality show | emotional clips |
|---|---|---|---|---|---|
| not included | 1st | **0.324 (0.007)** | 0.230 (0.084) | **0.363 (0.007)** | 0.254 (0.072) |
| | 2nd | −0.171 (0.131) | −0.154 (0.195) | −0.147 (0.195) | −0.158 (0.195) |
| included | 1st | **0.304 (0.014)** | 0.216 (0.070) | **0.336 (0.017)** | 0.239 (0.054) |
| | 2nd | **−0.265 (0.020)** | −0.198 (0.084) | **−0.276 (0.046)** | −0.247 (0.054) |

TABLE A4

Main results with age and sex not included as covariates (as reported in the manuscript) and with age and sex included as covariates (for comparison). AAL analysis. Statistically significant results ($q < 0.05$, FDR-corrected) are highlighted.

| covariates | PC | all | western | reality show | emotional clips |
|---|---|---|---|---|---|
| not included | 1st | **0.276 (0.026)** | 0.193 (0.253) | **0.297 (0.043)** | 0.154 (0.339) |
| | 2nd | −0.092 (0.413) | −0.069 (0.645) | −0.046 (0.685) | −0.081 (0.645) |
| included | 1st | **0.276 (0.027)** | 0.188 (0.262) | **0.299 (0.045)** | 0.157 (0.262) |
| | 2nd | −0.183 (0.107) | −0.114 (0.316) | −0.134 (0.285) | −0.154 (0.262) |

**Note:** Principal component (PC) orientation in PCA is arbitrary and may be reversed without affecting interpretation. For ease of comparison, we adjusted the orientation of the PCs in the ‘not removed’ analysis in Tables A1 and A2 so that they align with corresponding components in the ‘removed’ analysis for the results across all three videos (‘all’ analysis). This is a standard practice in PCA reporting to ensure consistent interpretation across conditions or datasets, and does not affect conclusions. Tables A3 and A4 did not require orientation adjustment, as the PCs were already aligned.
